# The temporal dynamics of the sea urchin regulome

**DOI:** 10.1101/2021.09.07.459266

**Authors:** Roberto Feuda

## Abstract

In this work, we used Nanostring N-counter technology, to evaluate the mRNA expression level of more than 330 regulatory genes over 34-time points covering the first three days of development of the sea urchin larvae. The hierarchical clustering of the mRNAs expression levels has identified groups corresponding to the major developmental landmarks (e.g., maternal to zygotic transition and gastrulation). Furthermore, comparison with previous experiments indicates high reproducibility of mRNA level temporal dynamics across batches. Finally, we generated an online tool to visualize gene expression during sea urchin larval development. The site can be accessed at https://nanostring2021.herokuapp.com/.

## Introduction

The larvae of sea urchin *Strongylocentrotus purpuratus* represent one of the best-studied organisms with a well-established Gene Regulatory Networks (GRN) up to early gastrula embryo (30 hours post fertilization – hpf- see for example(1–4)). However, several events crucial for the development of the larval body plan (e.g., the neuronal differentiation) happen in the post-gastrula stages where information on the mRNAs level is scant. There are about 280 transcription factors encoded in *Strongylocentrotus purpuratus* genome(5) and previous works have focused only on a few genes or at few developmental time points. Materna and colleagues analysed the mRNA expression level for 138 spatially restricted regulatory genes in early sea urchin development from fertilized egg to 48 hpf (6). Subsequently, Tu and collaborators sequenced the developmental transcriptome using mRNA-seq (7) for 10 timepoints but only three were from stages after 40 hpf. A comprehensive, high-density description of mRNA expression for the encoded transcription factors is crucial to identify genes involved in the establishment of the larval morphology. Furthermore, the dynamics in the mRNAs level can be used to infer potential regulatory interactions and the overall developmental hierarchy, the first critical step in the identification of the GRNs.

In this work, we measured the level of expression of 335 regulatory genes from zygote to 3 days of development, with at 2-hour interval for a total of 34-time points. We used Nanostring N-counter Technology that represents a middle-range technology where hundreds of genes can be measured simultaneously yet maintaining a high level of precision. Compared to other technologies, such as mRNA-seq and qPCR, the Nanostring N-counter Technology does not require a library or enzymes, reducing substantially the possible biases (8). In brief, the technology is based on the hybridization of two probes, a capture and reporter probe, to each target transcript and the number of hybridization events is quantified using an automated fluorescent microscope (8).

To study the mRNA temporal dynamics, we used hierarchical clustering and identified seven clusters that correlate with morphological events happening during sea urchin larval development). The data suggest that the level of expression of regulatory genes is sufficient to discriminate between developmental stages. Next, we compared our data with the previous observation from Materna and collaborators (6), and identified a high level of reproducibility for the mRNA expression level. Finally, to allow accessibility to the data, we developed an online tool that can be used to visualize the mRNA level of expression. The developmental time course can be found in a searchable database that is accessible at https://nanostring2021.herokuapp.com/.

## Materials and methods

### Code set

We designed a probe set containing 335 genes covering most of the regulatory genes expressed during the development of *Strongylocentrotus purpuratus* (see table S1 for the sequences). This includes transcription factors and other modulators of gene expression such as WNT and FGF signalling and a transcriptional cofactor (see table S2). The code set was designed using previous information from Materna et al. (6) and using gene models predicted by Tu et al. (7). The genes were classified using EggNog Mapper (9), and BLASTP against the Metazoan transcription factor database (10) (see Table S2 for the classification).

### Embryo culture and RNA extraction

Sea urchin embryos were fertilized in filtered seawater and cultured at 15°C. Every other hour ~300 embryos were counted and lysed in 350μl of a solution of RLT buffer and β-mercaptoethanol from the Qiagen RNeasy Micro Kit (Qiagen, Hilden, Germany). The lysates were immediately stored at −80°C until use. The RNA was extracted according to the manufacturer’s instructions but, similarly to Materna et al. (6), to maximize recovery, RNA was eluted with 100 μl nuclease-free water at 60°C. The samples were ethanol precipitated and resuspended in 7μl nuclease-free water, 5 of which were used in the following NanoString hybridization.

### Nanostring nCounter assays

For each individual time point the transcript count was measured using the NanoString nCounter. Hybridization reactions were performed according to the manufacturer’s instructions with 5μl RNA solution. Care was taken to minimize the time after the addition of the capture probe set to minimize background due to non-specific interactions between detection probes and capture probes. All hybridization reactions were incubated at 65°C for a minimum of 18 h. Hybridized probes were recovered with the NanoString Prep Station and immediately evaluated with the NanoString nCounter. For each reaction, 600 fields of view were counted.

### mRNA quantification

We performed two biological replicates, and the mRNA level was quantified as follows. First, for each sample, the background correction was performed individually by removing the total sum negative control using NanoStringNorm (11). This step removes aspecific probe binding. Second, to account for differences in hybridization efficiency, we estimated the mRNA counts using the Nanostring internal spikes. We took advantage of the six Nanostring exogenous mRNA spikes-in those covers from 0.125 femtomolar to 125 femtomolar. We converted this into mRNA molecules and estimated the mRNA level for each sample by comparing gene counts and mRNA spikes-in counts. While these steps ameliorate the differences in the hybridization events, they do not correct the differences between samples (e.g., different number of embryos) that can affect the reconstruction of the time-course. We combined the correct counts for all time points and used the total counts to account for the variation between time points. Finally, the average between the mRNA was used to combine the two replicates.

## Results and discussion

### Developmental time explains the majority of the observed variation

To quantify the variation between the two biological replicates, we performed a PCA using ClusVis (12).The results suggest that the majority of variance (63.8%) is explained by developmental time (Figure 1). In all 34 points, the two biological replicates are clustered together, suggesting a high level of similarity between replicates.

**Figure 1.**
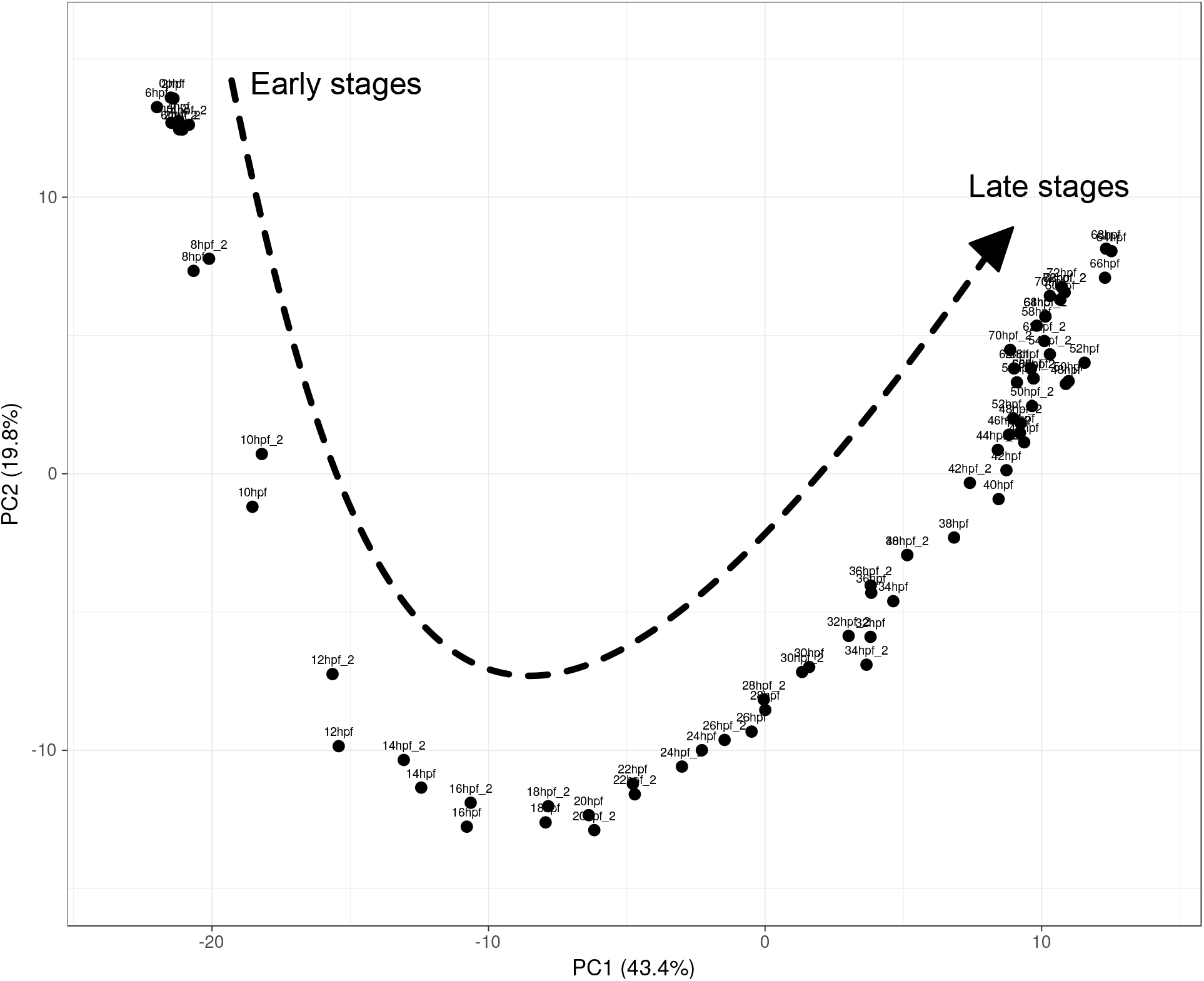
Principal component analysis of the temporal expression data. PCA analysis of Nanostring dataset from 0 to 72 hpf (*n* = 2 replicates per time point). The 63.8% of variance is explained by the gene expression temporal dynamics.

To clarify the temporal dynamics in gene expression, we performed a clustering analysis using ClusVis (12). The results in Figure 2 indicate the existence of multiple distinct clusters that correspond to well defined developmental stages defined based on embryological features. The first cluster confirms the results of the PCA and identifies a cluster that includes all the samples from 0 to 8 hpf that are consistent with previous data from transcriptional kinetics and suggest that the activation of the zygotic genome happens after 5-6 hours post fertilization (13). The remaining time points are organized in six distinct clusters that reflect the morphological events happening during sea urchin larval development. For example, one cluster comprises 10 to 18 hpf that encompass the majority of the Blastula stage. A similar pattern is observed for later stages where our clustering analysis identifies a sample cluster from 20 hpf to 26 hpf representing the mesenchyme blastula. These results indicate the expression of regulatory genes contains enough statistical power to identify well defined developmental stages. Furthermore, these results indicate that classic morphological changes happening during sea urchin development are the results of the changes in underlying expression of regulatory genes.

**Figure 2.**
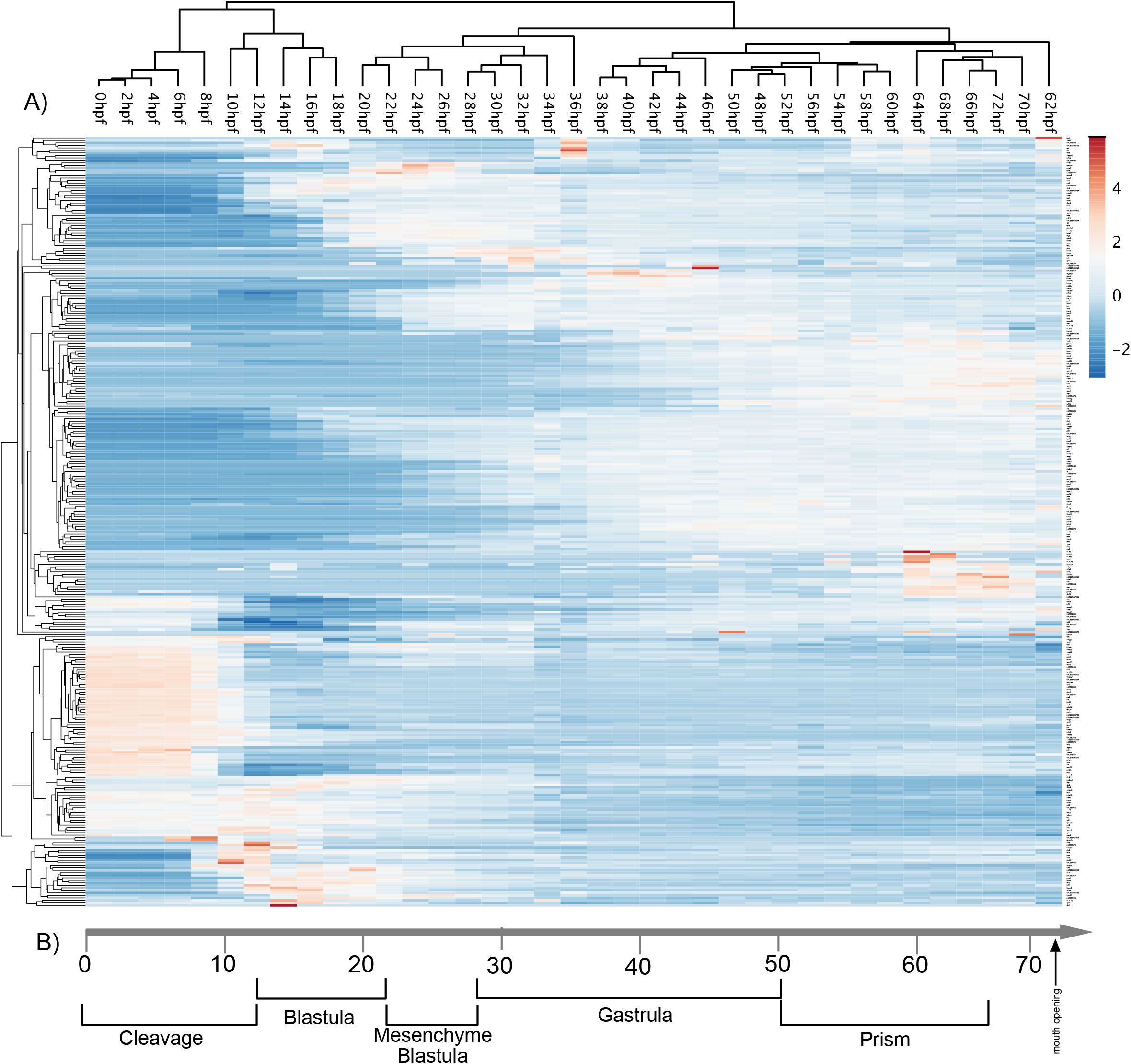
Heat map of gene expression. **(A)**Heat map of scaled expression for average mRNA expression level for 335 genes average obtained from two replicates from 0 to 72 hpf (34-time points) of sea urchin larval development. The results indicate the existence of the X clusters that reflect the developmental transition during sea urchin embryonic development (lower part of the figure). **(B)**Developmental stages of *S. purpuratus* embryos at 15°C

### High reproducibility of temporal dynamics of regulatory gene expression

Finally, we tested the reproducibility of our developmental time course. To this aim, we compared our expression profile with Nanostring data obtained by Materna and collaborators (6). In this case, the authors evaluated the level of expression to sea urchin development for 138 regulatory genes for the 48 hours of development. Differently from this work, Materna and collaborators used spike-GFP and RFP to normalize the data. To compare the datasets, we selected eight genes that have a different level of expression from 100 to 20000 copies of mRNA per embryo. Comparison (Figure 3) shows that results are highly consistent and reproducible, despite using different instruments, conditions, and animal batches. This indicates that the developmental progression of genome regulation is tightly controlled.

**Figure 3.**
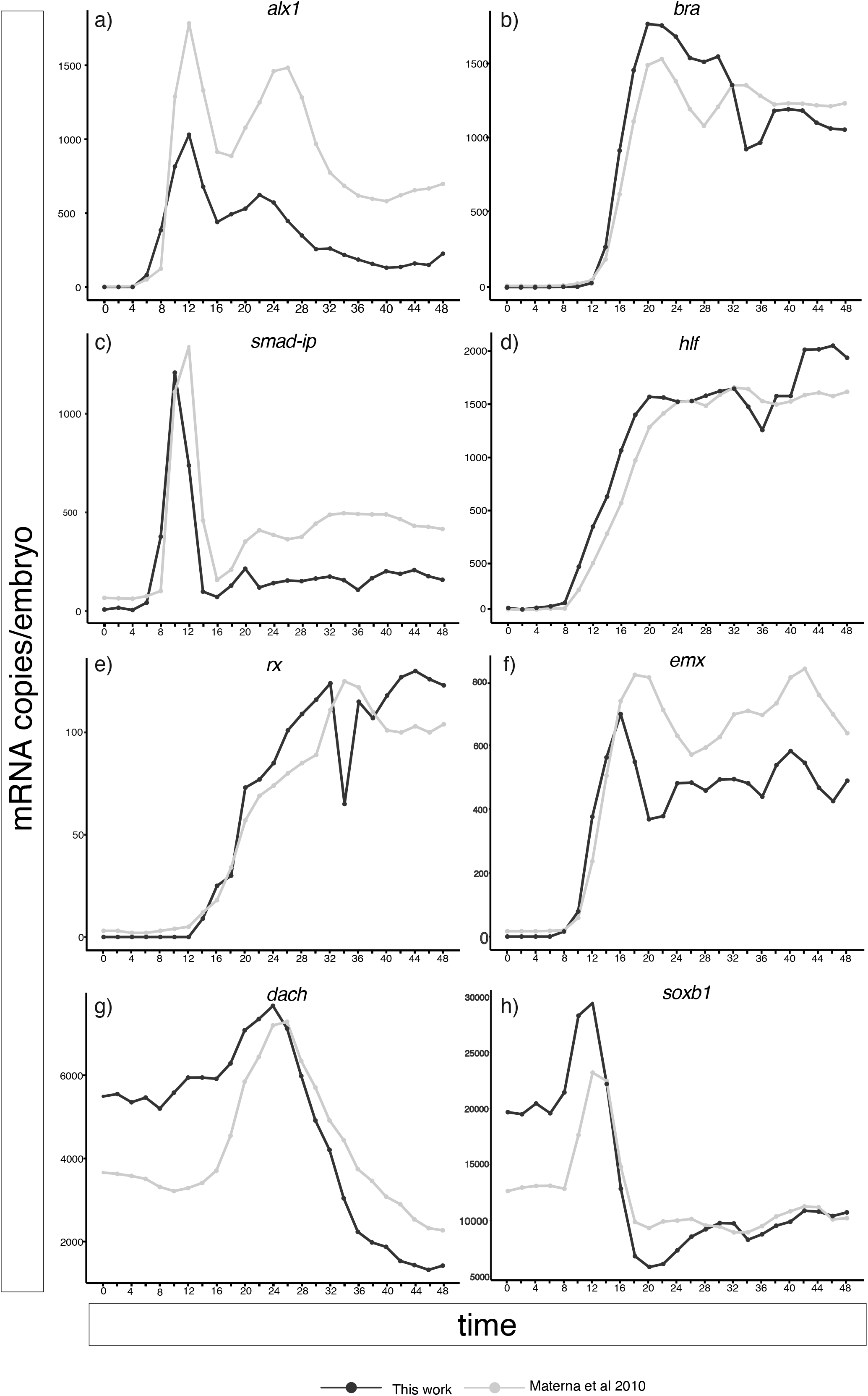
Reproducibility of mRNA measurements. Line plots obtained comparing mRNA expression from this work and those from Materna and collaborators (6). The results indicate a high level of reproducibility of Nanostring measurements as well as of the transcriptional hierarchy.

### Data accessibility

To visualize the mRNA expression for all the genes, we have created a visualization tool available at https://nanostring2021.herokuapp.com/. A screenshot is shown in Figure 4. With this tool, it is possible visualize the mRNA expression level for the 335 regulatory genes analysed in this work. A table with all expression data is available as supplementary data.

**Figure 4.**
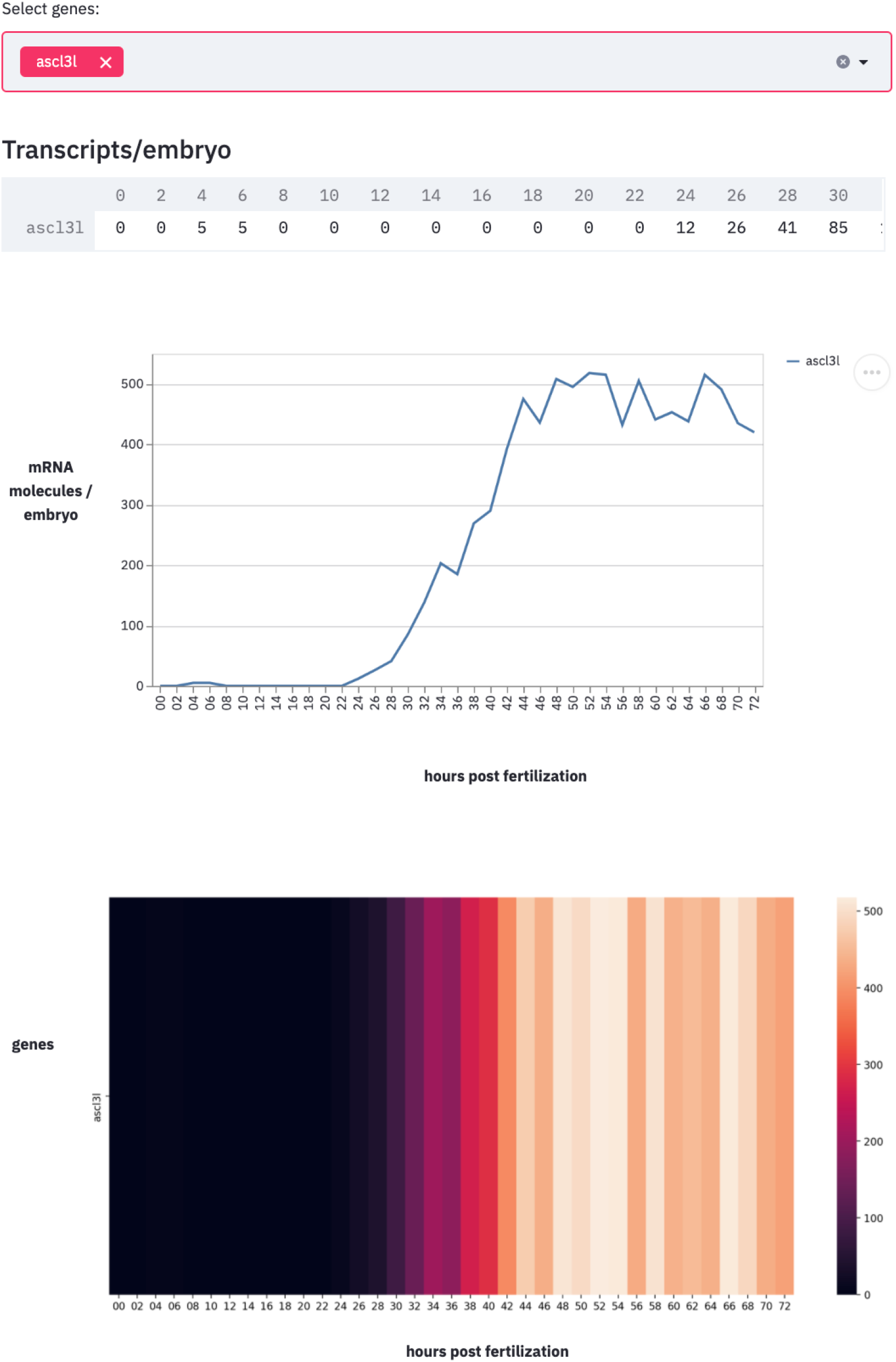
Snapshot of the expression data. The webpage allows for the visualization of up to 100 genes simultaneously both using heatmap or line plot.

### Conclusions

By describing the developmental dynamics, the data presented in this work might inform future functional perturbations. The high-density sampling and the inclusion of potentially all the known transcription factors data can be used to reconstruct the temporal developmental hierarchy between the regulatory genes, and together with spatial data, inform the functional perturbation

---

**Table S1. Nanostring codeset used in this work.** This table shows the gene names, WHL model and probe sequence for the 335 genes used in this work

**Table S2. Classification for the 335 regulatory genes used in this work.** This table shows the annotation for each of the 335 genes obtained using BLASTP and the Animal Transcription factor database and using EggNog Mapper.

**Table S3. Expression level for 355 regulatory genes generated in this work**. This table shows the expression level for 335 and 34 time points generated in this work.

## Acknowledgments

I would like to thank Eric Davidson and Isabelle Peter for their tireless support. Johnathan Valencia and Enhu Li for the help in embryos collections. Julius Barsi for designing the Codeset and Maryam Ghaffary Saadat for developing the visualization tool. Ezio Rosato, Alessandra Perugini and the Feuda lab for comments on the MS. This work is supported by a Royal Society University Research Fellowship (UF160226) and a Royal Society Research grant (RGF\EA\201005) to R.F.

## Notes

### Competing Interest Statement

The authors have declared no competing interest.

## References

1. P. Oliveri, Q. Tu, E. H. Davidson, Global regulatory logic for specification of an embryonic cell lineage. PNAS 105, 5955–5962 (2008).

2. I. S. Peter, E. H. Davidson, A gene regulatory network controlling the embryonic specification of endoderm. Nature 474, 635–639 (2011).

3. I. Peter, E. H. Davidson, Genomic control process: development and evolution (Academic Press, 2015).

4. E. Li, M. Cui, I. S. Peter, E. H. Davidson, Encoding regulatory state boundaries in the pregastrular oral ectoderm of the sea urchin embryo. PNAS 111, E906–E913 (2014).

5. M. Howard-Ashby, et al., High regulatory gene use in sea urchin embryogenesis: Implications for bilaterian development and evolution. Dev Biol 300, 27–34 (2006).

6. S. C. Materna, J. Nam, E. H. Davidson, High accuracy, high-resolution prevalence measurement for the majority of locally expressed regulatory genes in early sea urchin development. Gene Expr. Patterns 10, 177–184 (2010).

7. Q. Tu, R. A. Cameron, E. H. Davidson, Quantitative developmental transcriptomes of the sea urchin Strongylocentrotus purpuratus. Dev Biol 385, 160–7 (2014).

8. G. K. Geiss, et al., Direct multiplexed measurement of gene expression with color-coded probe pairs. Nature Biotechnology 26, 317–325 (2008).

9. J. Huerta-Cepas, et al., eggNOG 5.0: a hierarchical, functionally and phylogenetically annotated orthology resource based on 5090 organisms and 2502 viruses. Nucleic Acids Res. 47, D309–D314 (2019).

10. H. Hu, et al., AnimalTFDB 3.0: a comprehensive resource for annotation and prediction of animal transcription factors. Nucleic Acids Research 47, D33–D38 (2019).

11. D. Waggott, et al., NanoStringNorm: an extensible R package for the pre-processing of NanoString mRNA and miRNA data. Bioinformatics 28, 1546–1548 (2012).

12. T. Metsalu, J. Vilo, ClustVis: a web tool for visualizing clustering of multivariate data using Principal Component Analysis and heatmap. Nucleic Acids Res 43, W566–570 (2015).

13. T. Gildor, A. Malik, N. Sher, S. Ben-Tabou de-Leon, Mature maternal mRNAs are longer than zygotic ones and have complex degradation kinetics in sea urchin. Developmental Biology 414, 121–131 (2016).

